# Constructing and validating a transferable epidemic risk index in data scarce environments using open data: a case study for dengue in the Philippines

**DOI:** 10.1101/2021.02.23.432447

**Authors:** Fleur Hierink, Jacopo Margutti, Marc van den Homberg, Nicolas Ray

## Abstract

Epidemics are among the most costly and destructive natural hazards globally. To reduce the impacts of infectious disease outbreaks, the development of a risk index for infectious diseases can be effective, by shifting infectious disease control from emergency response to early detection and prevention.

In this study, we introduce a methodology to construct and validate an epidemic risk index using only open data, with a specific focus on scalability. The external validation of our risk index makes use of distance sampling to correct for underreporting of infections, which is often a major source of biases, based on geographical accessibility to health facilities. We apply this methodology to assess the risk of dengue in the Philippines.

The results show that the computed dengue risk correlates well with standard epidemiological metrics, i.e. dengue incidence (p = 0.002). Here, dengue risk constitutes of the two dimensions susceptibility and exposure. Susceptibility was particularly associated with dengue incidence (p = 0.047) and dengue case fatality rate (CFR) (p = 0.029). Exposure had lower correlations to dengue incidence (p = 0.211) and CFR (p = 0.163). Highest risk indices were seen in the south of the country, mainly among regions with relatively high susceptibility to dengue outbreaks.

Our findings reflect that the modelled epidemic risk index is a strong indication of sub-national dengue disease patterns and has therefore proven suitability for disease risk assessments in the absence of timely epidemiological data. The presented methodology enables the construction of a practical, evidence-based tool to support public health and humanitarian decision-making processes with simple, understandable metrics. The index overcomes the main limitations of existing indices in terms of construction and actionability.

**Author summary:**

- Why Was This Study Done?
  – Epidemics are among the most costly and destructive natural hazards occurring globally; currently, the response to epidemics is still focused on reaction rather than prevention or preparedness.
  – The development of an epidemic risk index can support identifying high-risk areas and can guide prioritization of preventive action and humanitarian response.
  – While several frameworks for epidemic risk assessment exist, they suffer from several limitations, which resulted in limited uptake by local health actors - such as governments and humanitarian relief workers - in their decision-making processes
- What Did the Researchers Do and Find?
  – In this study, we present a methodology to develop epidemic risk indices, which overcomes the major limitations of previous work: strict data requirements, insufficient geographical granularity, validation against epidemiological data.
  – We take as a case study dengue in the Philippines and develop an epidemic risk index; we correct dengue incidence for underreporting based on accessibility to healthcare and show that it correlates well with the risk index (Pearson correlation coefficient 0.69, p-value 0.002).
- What Do These Findings Mean?
  – Our methodology enables the development of disease-specific epidemic risk indices at a sub-national level, even in countries with limited data availability; these indices can guide local actors in programming prevention and response activities.
  – Our findings on the case study show that the epidemic risk index is a strong indicator of sub-national dengue disease patterns and is therefore suitable for disease risk assessments in the absence of timely and complete epidemiological data.

## 1 Introduction

### Epidemic Risk Assessment

Epidemics are among the most costly and destructive natural hazards globally [1, 2]. Currently humanitarian action to epidemics is focused on response rather than preparedness and prevention [3–6]. Timely detection of disease cases in combination with risk assessment can support prevention measures and therefore contribute to early containment of outbreaks [4, 6]. The use of a holistic risk index for infectious diseases can reduce the impacts of epidemics on (vulnerable) communities, by shifting infectious disease control from response after emergence to early detection and prevention [1, 2, 6]. Comparable risk indices for natural hazards and humanitarian crises have proven to be effective in localizing high risk regions [7, 8] and are being used to inform preparedness programs [7, 8]. Research has shown that the risk of re-emerging infectious disease outbreaks or new spillover events (i.e. pathogen transmission from a reservoir to a new host) is increasing due to degrading ecosystems, intensification of travel and trade, climate change, population growth, and a wide variety of other factors [3, 6]. It is therefore imperative to increase our understanding of disease risk distribution at the most local level possible, so impacts can be reduced accordingly [4–6].

Epidemic risk is usually quantified by several indicators, which relate both to the probability of outbreak occurrence and to its potential impact [9–12], chosen according to the specific disease(s) under consideration. These indicators are combined and mapped to a normalized risk index, according to initial estimates (most commonly, weighted or geometric means). They can be conceptually divided into two dimensions [9]:

1. Hazard and exposure: the presence of an infectious disease and its vector (e.g. mosquitoes) and the likelihood of exposure
2. Vulnerability and coping capacity: clinical, demographic and socioeconomic data that influence health outcomes (e.g. age); the ability of a government or health system to detect, contain and respond to an outbreak (e.g. hospital capacity)

While several frameworks for epidemic risk assessment exist [9–13], they have been hardly used by health actors - such as governments and humanitarian relief workers - to prioritize intervention areas and actions, despite them being the first responders to epidemic outbreaks and thus often carrying major decision responsibilities. While dedicated research on the reasons behind this lack of adoption is missing, anecdotal evidence from humanitarian practitioners often points to one or more of the following limitations, which existing epidemic risk indices suffer from:

1. Methodology:
  a. rely on accurate clinical, virological and/or entomological data, which often require dedicated and in-situ data collection campaigns; these are costly, impractical and often prerogative of health authorities
  b. focus on a global scale, by comparing world regions or countries; this can be useful for international organisations (e.g. in long-term planning of funding by donors), but not for local ones [14]
  c. lack of validation against epidemiological data
2. Actionability: lack a clear connection with with policy implications and practical interventions, i.e. a prescriptive aspect [15]

The methodological limitations are connected with data availability, most importantly of clinical surveillance data, whose lack of determines the difficulty, respectively, of using advanced epidemiological models [16], of modelling at a sub-national scale and, finally, of validating results. While epidemic surveillance systems that collect and aggregate this data [17] do exist, they often lack completeness and timeliness, especially in developing countries [18], which carry the highest burden of infectious diseases [1]. Official ures from such surveillance systems are often derived from clinical records of symptomatic cases [19], i.e. *passive surveillance*, and thus do not take into account asymptomatic cases, underdiagnosis and, most importantly for low- and middle-income countries (LMICs) countries, infected individuals who do not receive treatment. This problem is often referred to as underreporting.

Underreporting in passive surveillance systems has been recognised as a major source of bias in estimates of infectious disease incidence, especially in LMICs [20, 21]. Known factors related to underreporting, other than possible asymptomatic cases [22], are sociodemographic factors that impede health-seeking behavior, such as poverty and education [23, 24], and geographical accessibility to health facilities [25–27]. Understanding and quantifying these factors is thus necessary to assess disease incidence, and thus epidemic risk, at a local level [26]. While the socioeconomic indicators which affect self-reporting are usually difficult to measure at a population-level and their relative importance is highly dependent on the local context [23], data on the location of health facilities is available virtually world-wide (but with various degrees of completeness) and their geographical accessibility can be modeled with suitable geo-spatial tools [25, 28]. Geographical accessibility models present an important and novel opportunity to bridge the data gap between reported and unreported cases, as it reflects the ability of a population to reach a health facility within a certain travel time [28]. Recently, a robust methodology has been proposed to correct for underreporting based on known covariates [26].

### Dengue and the Philippines

Vector-borne diseases (VBDs), i.e. infectious diseases that are transmitted through a blood-feeding arthropod (e.g. mosquitoes, sandflies, ticks, etc.) are an important group of infectious diseases [29]. VBDs are responsible for 17% of the total burden of all communicable diseases and their prevalence disproportionately affects the poorest communities in tropical- and subtropical regions [29, 30]. Socioeconomic, demographic and environmental indicators are known to be strongly linked to the distribution of VBD risk and an expansion of transmission patterns in the coming years is expected due to environmental changes, rapid urbanization, and globalization [30].

While the global communicable disease burden for some of the largest infectious diseases (i.e. HIV/Aids, tuberculosis, and malaria) has been tremendously reduced over the last decade, deaths due to the VBD dengue have increased by 65.5% from 2007 to 2017, with the same trend seen for dengue case fatality rates (CFR) [29, 31]. Dengue is a mosquito-borne disease with four different serotypes (i.e. DENV-1, DENV-2, DENV-3, DENV-4) and is considered as a neglected tropical disease by the World Health Organization (WHO). The disease is spread by the mosquitoes *Aedes aegypti* and *Aedes albopictus* and is responsible for an estimated 96 million cases annually, with 50% of the world’s population expected to be at risk [29, 30]. Currently, the primary method for controlling dengue are vector control strategies, aimed at limiting human exposure to the transmitting mosquitoes. Targeting regions for vector control measures is of high importance to optimize and maximize the effect of the available resources [29, 30]. Understanding the distribution of dengue risk is key in tailoring and targeting intervention strategies on sub-national scales [30], but challenging due to underreporting [26]. Improved methods are needed to meet the recently updated WHO NTD roadmap target of a 0% dengue case facitliy rate (CFR) by 2030 (from the 0.8% baseling in 2020) [32].

Dengue is a large scale health challenge in The Philippines [33]. The disease is endemic in the entire country with re-occurring outbreaks in all regions and the circulation of all virus strains. The country is highly vulnerable for dengue outbreaks, partly as a consequence of recurring natural hazards destructing critical infrastructures, but also because of environmental conditions favouring the life-cycle of mosquitoes [34, 35]. Dengue surveillance in the Philippines mostly represents hospitalized cases, particularly those of patients with severe dengue infections. Between 2010 and 2014 about 93% of all reported dengue cases concerned hospitalized patients of which 50% were reported from private facilities [33]. This finding highlights the fact that a large portion of the dengue cases may remain unreported, hindering a realistic understanding of dengue in the country and thus stressing the need for realistic correction methods [33].

Reliable risk estimates of dengue are needed in the Philippines to allow guided allocation of preventive measures and targeted outbreak containment [33, 36]. Research on dengue in the Philippines has focused on modeling techniques, with the goal of either describing past disease dynamics or to predict future ones [34, 37–40]. While such models could provide estimates of (future) morbidity, which is a key ure to inform epidemic response and preparedness programs, they suffer one or more of the following limitations: relying on detailed clinical and/or entomological data, which is rarely available, and not discussing potential health outcomes, e.g. by considering the local (health) capacity. To the best of our knowledge, no research has been carried out yet on combining the different dimensions of dengue risk (hazard and exposure, vulnerability and coping capacity) into one quantitative framework, which can be applicable at a country scale. The inherent challenge, like in other data-scarce environments, is missing information on relevant risk indicators and epidemiological data.

In this study, we present a methodology to build and validate an epidemic risk index at a sub-national level, using openly available data and tools, to ensure its applicability in data-scarce settings. The methodology can be conceptually divided in three steps:

- Development of the Epidemic Risk Index: selection of indicators, normalization and aggregation
- Correction of (public) epidemiological data: estimation of relative differences in underreporting based on geographical accessibility to healthcare
- Validation of the Epidemic Risk Index against corrected epidemiological data

## Materials and methods

### Study region

The Philippines is an archipelago nation in the Western Pacific ocean and is subdivided into 17 administrative regions, which are further subdivided into 81 provinces, 1489 municipalities and 42,036 barangays [41, 42]. Historically, dengue cases were reported on a weekly basis by the Department of Health in surveillance reports [43]. Although we acknowledge that spatial granularity is important in risk assessment models, dengue cases have been mostly openly reported on regional level (n = 17). Therefore this study focuses on a risk index for all 17 administrative regions in the Philippines.

### Epidemic Risk Index

The epidemic risk index was built largely following the methodology of the Water Associated Disease Index (WADI) [12], which has been developed with a specific focus on dengue and has been successfully validated against actual dengue incidence data, even at a sub-national level, in Malaysia [13] and Vietnam [44]. The risk index is defined as a weighted average of two components, exposure and susceptibility, which quantify the risk of being exposed to the pathogen and the risk of experiencing severe health outcomes, respectively. Following the methodology of [13], the weights in this average were chosen to maximize the correlation with dengue incidence.

Each component is in turn defined as the arithmetic average of one or more indicators, summarized in Tab. 1. Each indicator, unless already normalized so that it lies in the range [0,1], was transformed according to

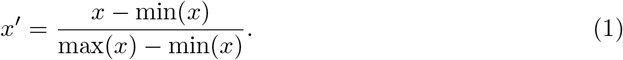

Concerning the choice of indicators, exposure was quantified as the fraction of the population *E* exposed to *Aedes aegypti*. This indicator was calculated from the probability of occurrence of the main dengue vector (*Aedes aegypti*) *V*, modeled in a raster format [45], and the population density distribution *ρ* [46], according to

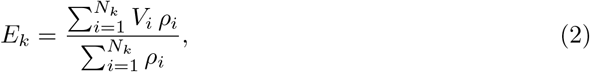

where *N_k_* is the number of raster cells within the boundaries of region *k*. We used high resolution population density estimates from the Facebook Connectivity Lab and Center for International Earth Science Information Network (CIESIN) [46].

Susceptibility was instead quantified combining five indicators which relate to vulnerabilities against dengue. Children 0-15 years of age have much higher chances to develop severe dengue with respect to the adult population [47] and thus constitute the most vulnerable group; their relative abundance was quantified via the fraction of the population belonging to the corresponding age group. Secondly, education has been identified among the key factors enabling health-seeking behavior [23], especially for the caregiver of the household [48], and is assumed to increase the capacity of interpreting and acting upon public health information aimed at preventing dengue. This was quantified via the female enrollment ratio to secondary school. Thirdly, the percentage of households using unimproved sanitation facilities (pit latrines without slabs or platforms or open pit, hanging latrines, bucket latrines, open defecation) was used to capture the risk of having exposed water containers in the house, which can act as breeding sites and was associated with higher risk of dengue [49]. Both this and the second indicator (education) are effectively proxies for poverty, which indirectly affects health outcomes [50]. Lastly, the density of physicians and public hospital beds was used to quantify, respectively, the availability and affordability of healthcare Table 1.

**Table 1.** Indicators used to build the Epidemic Risk Index and epidemiological metrics to validate it.

| Component | Indicator | Rationale for inclusion | Data source |
| --- | --- | --- | --- |
| Exposure | Fraction of the population exposed to *Aedes aegypti* | *Aedes aegypti* is the main dengue vector in the Philippines | [45, 46] |
| Susceptibility | Percentage of children 0-15 years of age | Children 0-15 years of age have higher susceptibility to severe dengue and CFR is higher among this group | [54] |
|  | Female enrollment ratio to secondary school | Progression to secondary school indicates a sufficient level of education and attainment to read, interpret and act upon public health information about dengue | [55] |
|  | Percentage of households using unimproved sanitation facilities | Unimproved sanitation indicates poorly managed water resources and poor housing quality | [56] |
|  | Number of physicians per 1000 people | Physician density is a proxy for availability of healthcare | [57] |
|  | Number of beds in public hospitals per 1000 people | Density of beds in public hospitals is a proxy for affordability of healthcare | [58] |
| Validation | Dengue incidence (number of dengue cases per person per year) and CFR; both were averaged over the years 2014-2019. | Average incidence and CFR quantify, respectively, the risk of outbreaks and their severity, in terms of health outcomes | [43] |

While geographical accessibility to health facilities is equally important in determining the probability of seeking and receiving adequate treatment [51], it was not included in the definition of our risk index to avoid a spurious correlation with the dengue incidence, which was corrected using accessibility data and was ultimately used for validating the index. The risk of each dengue case to develop into severe dengue and, potentially, death is known to be determined by a number of other factors [52], most importantly immunity and previous exposure to a different serotype of dengue, due to antibody-dependent enhancement [53]; however, to the best of our knowledge, no serotype-specific case data exists for the region and period under study and we thus had no way to quantify such effect. Ultimately, the calculated risk index was validated against the dengue CFR and incidence [43], averaged over the years 2014-2019, by means of Pearson correlation coefficients.

#### Accessibility to health care

Accessibility to health care was measured in terms of travel time (in minutes) to the closest health facility with dengue testing services. The applied travel scenario considered motorized travel speeds on roads and walking travel speeds on other land cover types (e.g. forest, grassland, urban landscapes) under the assumption that patients walk to the nearest road and then continue their journey with a vehicle that is readily available. The travel time raster was modelled by means of a least cost-distance algorithm in AccessMod version 5.6.30 [28]. In order to obtain a single 110 meter resolution raster travel impedance surface, spatial data on elevation, land cover, roads, and river networks were merged in an overarching raster layer through the*merge landcover* module in AccessMod, to which the travel scenario was applied [28] S1.

Data preparation of all separate spatial layers was done using RStudio (R version 4.0.2). Land cover data was downloaded in tiles from Coopernicus [59] and elevation data from Shuttle Radar Topography Mission (SRTM) [60]. Both spatial raster layers were mosaiced to cover the Philippines and clipped to country borders. The two raster layers (land cover and elevation) were then re-sampled to a resolution of 110 meter, using the native resolution of the landcover as a reference, and raster cells were aligned with the elevation layer as a reference.

Vector data representing the road network and hydrography had to be separately downloaded for the Northern and Southern part of the country from Humanitarian Open Street Map [61, 62] and was enriched with data from the Open Mapping at Facebook Initiative [63]. Layers on both parts of the country were merged. Hydrographic features such as rivers and lakes were considered full barriers to movement to the population, unless a road crosses over, which was considered as a functional bridge. Road data was cleaned to only contain OpenStreetMap official road classes [64] and new integer road class values were created for each unique road type, as an essential step for the land cover merge. Health facility coordinates were downloaded from the Department of Health in the Philippines [65] and health facilities known to offer dengue testing services (i.e. “Rural Health Unit”, “Hospital”, “Medical Clinic”), as discussed with country representatives were filtered from the data. Coordinates falling on barriers were moved to the nearest neighbouring non-barrier cell and facilities wrongly located far outside country borders were removed from the analysis. All raster and vector datasets were projected to the Philippines’ projection system (EPSG:32651, UTM51N).

#### Reporting probability

The travel time raster obtained from the accessibility model served as the input data for the calculation of the reporting probability, following a distance sampling methodology [26]. In particular, we used the following equation to describe the reporting probability (*P*) as a function of travel time to the nearest health facility with dengue testing service (*t*):

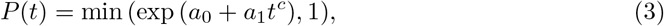

where *a*_0_, *a*_1_ and *c* are free parameters. This function captures the main feature of the traditional assumption used in distance sampling methods, namely an exponential decrease. Since we did not have access to individual patient case data, we used the results of [26] to give an estimate of the free parameters in Eq. 3, converting the time travel *t* to distance *d* by dividing it by the average travel time *v*:

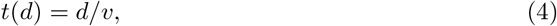

where *v* was estimated according to the aforementioned travel scenario (S1).

Eq. 3 was first applied to each cell of the travel time raster, to produce a reporting probability raster. We then computed the average reporting probability per region 〈*P*〉 by taking a weighted average of the reporting probability within the region boundaries and using population density *ρ* as weight:

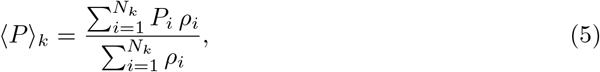

where *N_k_* is the number of cells within the boundaries of region *k*. We used the same population density estimates [46], resampled to 110m resolution by summing population. This technique results in the loss of population across the grid, mainly as a result of reprojecting the layer. To correct for this, the total lost population was smoothed out over the resampled population grid.

Average reporting probability was then used to correct dengue regional incidence, which was in turn estimated from official dengue case counts [43] and census data [54]. This step corrects for the major imbalance in official dengue statistics due to unequal access to healthcare. Finally, since the almost entirety of reported cases comes from hospitalized settings [66] due to the dengue case definition [67], dengue incidence was corrected for the fraction of hospital beds belonging to facilities connected to the Philippines epidemiological surveillance system [68].

## Results

### Validation of the Epidemic Risk Index

The Epidemic Risk Index and its components are validated against the corrected dengue incidence and CFR in the 17 regions under study by measuring the Pearson correlation coefficient *r* (Table 2). The significance of each correlation is measured with *p*-value at the significance level of 0.05 (*p* < 0.05). Concerning incidence, a positive, significant correlation is observed between incidence and susceptibility and between incidence and risk: *r* = 0.49 (*p* = 0.047) and *r* = 0.69 (*p* = 0.002), respectively. Concerning CFR, a significant correlation is observed only between CFR and susceptibility, with *r* = 0.53 (*p* = 0.029).

**Table 2.** Correlation coefficients between the corrected dengue incidence, CFR, Epidemic Risk Index and its components. In bold: significant correlations (*p* < 0.05).

| Variables | Pearson $r$ | $p$-value |
| --- | --- | --- |
| Incidence and exposure | 0.32 | 0.211 |
| **Incidence and susceptibility** | **0.49** | **0.047** |
| **Incidence and risk** | **0.69** | **0.002** |
| CFR and exposure | -0.35 | 0.163 |
| **CFR and susceptibility** | **0.53** | **0.029** |
| CFR and risk | 0.25 | 0.342 |
| Exposure and risk | 0.44 | 0.081 |
| **Susceptibility and risk** | **0.73** | **0.001** |

#### Accessibility to healthcare

Accessibility to dengue reporting facilities was highest in the National Capital Region (Fig 1A-B). Where 99.98% percent of the population (N = 12,304,651) was able to reach a health facility within 1 hour travel time. Lowest accessibility coverage was seen in Region IV-B, with 83.3% percent of the population being able to access care in 1 hour (Fig 1A-B).

**Fig 1.**
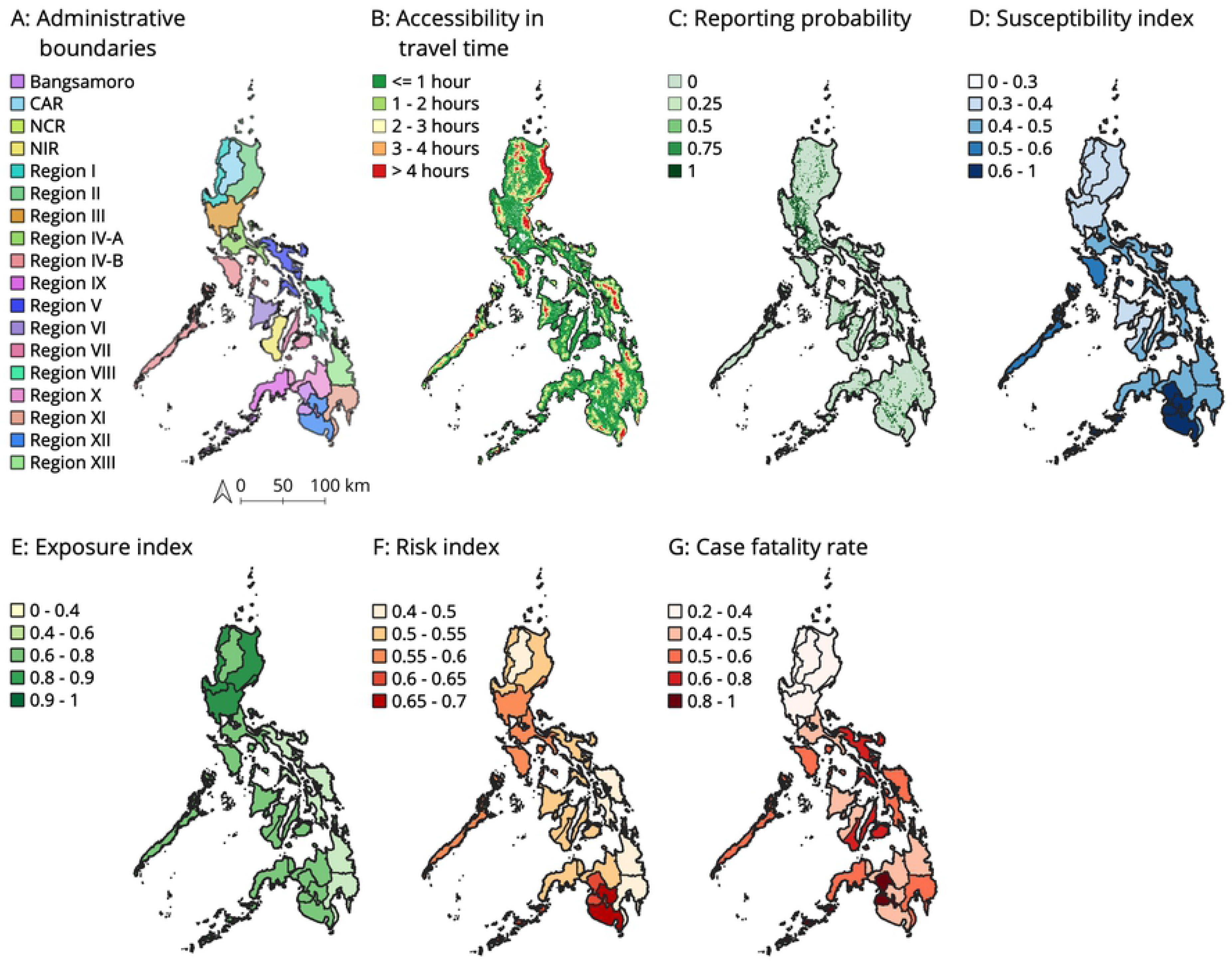
Overview of all results. Panel A shows the 17 administrative regions of the Philippines. Panels B-G highlight the individual results of the accessibility analysis, reporting probability, the dimensions that compose the risk index, the risk index, and the case fatality rate. Enlarged versions of the figures can be found in S3-S9

#### Reporting probability

Reporting probability (Fig 1C) was generally highest around Manilla with an average reporting probability of 0.89 in the National Capital Region. However, reporting probability was much lower for all other regions, with probabilities ranging from 0.10-0.37, implying that reported incidence was corrected with higher correction factors among all these regions (S5 and S10).

#### Geographical distribution of dengue risk

The maps in Fig 1D-F and S 1 represent the results of the calculated susceptibility, exposure, and ultimately risk index. Regions with high susceptibility, thus reflecting low coping capacity and resilience, are depicted in darker blue colors. Regions with high potential *Aedes aegypti* exposure are shown in darker green. Ultimately, regions with a relatively high risk index (Fig 1F) are shown in darker orange.

The Pearson correlation was strongest between the susceptibility dimension and dengue incidence (P = 0.047), as compared to the other covariates (Table 2). Therefore, susceptibility related variables weighted heavier on the risk index than the exposure variables. Susceptibility was highest in Region XII (0.65) and lowest in the National Capital Region (0.29), reflecting higher coping capacity of individuals and the health system around the capital.

Comparing the exposure index to the susceptibility index for instance, shows that regions with highest exposure index are mainly located in Northern regions of the country, whereas susceptibility was found to be highest in more Southern regions. The exposure index was found to be highest in the National Capital Region (0.95) and ranged from 0.43-0.95 throughout the entire country.

The modelled risk index ranged from 0.43 to 0.68 between all regions in the Philippines. All results are aggregated on regional level, firstly because dengue data was richest in terms of temporality and secondly because decision-making on resource allocation is often carried out at this level. The modelled risk index was highest for Region XII, with an index of 0.68 and the exposure and susceptibility index being 0.76 and 0.65 respectively (S2). Interestingly, CFR in this region was relatively low, being 0.42. The second highest risk index was seen in Bangsamoro, with a risk index of 0.63, and an exposure and susceptibility index of 0.66 and 0.62. In general, a cluster of higher risk indices was concentrated in Southwest Philippines (Fig 1F). When comparing this cluster of high risk indices against the susceptibility and exposure index, it becomes apparent that especially the susceptibility index is highest in these regions (Fig 1D) while higher values for the exposure index are seen among northern regions in the Philippines (Fig 1E). Highest CFRs are also concentrated in the Southwest regions of the Philippines (Fig 1G).

In general, when comparing the spatial distribution of the susceptibility and exposure index against the risk index there is no notable trend visible between the exposure index and the risk index. Yet, the susceptibility and risk indices show a more closely related trend, towards the southern regions of the country.

While accessibility in terms of travel time (Fig 1B) are highest in the Northeast of the Philippines, which might potentially reflect a poorer capacity to deal with an outbreak, the susceptibility index is generally low in this region.

## Discussion

In this study, we have constructed and validated an epidemiological risk index using openly available data, to assess exposure and vulnerability to dengue in the Philippines. The proposed methodology can be easily applied to other countries and diseases, as it does not use data which is uniquely available in the Philippines nor it depends on specific features of dengue epidemiology. More specifically, the indicators used to construct the index are commonly captured at a sub-national level by public demographic and health surveys or, where government capacity is limited, by humanitarian programs such as USAID’s DHS [69]; while different indicators might be more suitable for different diseases (e.g. elderly, not children, might be more at risk of severe health outcomes), we think that a reasonable set can be found within the aforementioned sources. The correction procedure of official epidemiological data for underreporting, which was used to validate the epidemic risk index, is also expected to be applicable in other contexts, i.e. other endemic infectious diseases and countries in which a passive surveillance system is in place.

Looking at our particular case study, we have shown how risk factors of dengue vary within the Philippines and how these correlate with epidemiological metrics. We observed, overall, that the combination of exposure and susceptibility explains, to some extent, the observed incidence and mortality rate, and it does so better than considering each of these two separately. The higher correlation between dengue incidence and risk index with regard to exposure and susceptibility alone is consistent with the hypothesis that there is an interplay between the latter two and that both need to be taken into account to correctly estimate epidemiological risk.

We also acknowledge that our study dealt with several challenges, which we discuss more in detail in the following.

### Accessibility analysis

Our travel scenario may not have been representative of all populations in the Philippines. Regional specificities on modes of travel or road quality may exist, and socio-economic differences within or between regions may alter the predominant modes and speeds of travel. A finer grain study on these potential geographic disparities could improve our travel model and therefore the reporting bias estimates.

### Correction for underreporting

While the models in [26] were fitted on case data of malaria in Burkina Faso, we argue that such scenario should be reasonably representative of dengue in the Philippines, at least for the purpose of this work. Dengue and malaria share indeed a high prevalence of asymptomatic cases [70, 71] which do not prompt healthcare seeking; also, they are both endemic in the Philippines and Burkina Faso, respectively. Other factors influencing health-seeking behavior, such as socio-economic ones [23], could determine a difference in reporting probability between these two countries; however, the factors that were explicitly included in [26] determined a poorer model performance with respect to including only distance, suggesting that the latter is indeed the main driver behind reporting probability. Finally, we note that our methodology aims at correcting for relative differences in reporting probability among regions in the Philippines, meaning that an absolute difference with true reporting probability might very well exist and does not influence the validity of our results.

### Risk index

The risk index that we constructed is meant to be a simple metric to guide decision-making processes and resource allocation of humanitarian agencies. Simplicity comes at a price: while we show that it does correlate with both incidence and CFR, and present a methodology to test this case-by-case, it is difficult to be more specific about actual expected health outcomes in case of an outbreak, given a certain value of the risk index.

Concerning exposure, the probability of vector occurrence has been modeled on the basis of environmental variables, among which the degree of urbanization (urbanicity) [45]; however, such model did not explicitly take into account the abundance of breeding sites, most importantly in solid waste and plastic containers, which has recently been identified as a key ingredient of vector ecology [72, 73]. The type and coverage of solid waste management is therefore expected to be a good predictor of vector abundance, although geographically detailed information on such a topic in the study region is scarcely available. Also, we note that using climatic averages to compute vector exposure is another important limitation of [45], as dengue incidence is known to follow seasonal patterns in the Philippines [35, 37]. However, extensive research as been conducted already on the topic [34, 40, 74] and a time-dependent exposure is easily implementable within the current framework, enabling real-time monitoring or even forecasts of the risk index throughout the year. We plan to address this in future research.

Finally the susceptibility dimension does not hold information on potential transmission dynamics of dengue to the population it represents the social predisposition and resilience of the population in case an outbreak occurs. Therefore, it can help target regions for building prevention and preparedness strategies. While susceptibility indicators capture important aspects of health systems, they might not take into account local, specific factors that play a decisive role both in health-seeking behavior and capacity to deliver care. In the Philippines, for instance, the southwest region of Bangsamoro (previously known as Autonomous Region in Muslim Mindanao) has been plagued by years of violent conflict between tribal, political and religious group and the government [75]. Not only does this affect the local health system resilience, but it is also a major factor to consider when planning humanitarian interventions, which this risk index is meant to inform.

## Conclusions

The presented methodology enables the construction of a practical, evidence-based tool to support public health and humanitarian decision-making processes with simple, understandable metrics, namely the epidemic risk index and its components. Our methodology overcomes the main limitations of existing epidemic risk indices (see Introduction): it is based on openly available data, it is localized, and results can be validated against epidemiological data. In terms of actionability, other than helping prioritizing intervention areas, we note that individual indicators contain useful information for humanitarian programs. Absolute numbers of potentially exposed and vulnerable people, for instance, can be directly extracted, together with clear indications on which interventions should be prioritized and where (e.g. vector control programs versus strengthening community-based surveillance). Investments in epidemic prevention, detection, and response are needed to advance in our capacity to deal with infectious disease outbreaks. The information captured in the epidemic risk index supports the general shift from reaction after emergence to epidemic prevention and preparedness that has been so widely advocated for, especially in light of the ongoing COVID-19 pandemic, and is transferable to other infectious diseases and settings.

## Supporting information

**S1 Tab. Travel scenario.** The speeds applied to our friction raster to calculate the accumulated travel times to dengue reporting facilities.

**S2 Fig. Overview of gradient indices.** Absolute indices for individual dimensions, risk index and case fatality ratio.

**S3 Fig. Regional boundaries in the Philippines.**

**S4 Fig. Accessibility to dengue reporting facilities expressed as travel time.**

**S5 Fig. Reporting probability of dengue.**

**S6 Fig. Susceptibility index.**

**S7 Fig. Exposure index.**

**S8 Fig. Risk index.**

**S9 Fig. Case fatality ratio.**

**S10 Tab. Correction factors.**

## Acknowledgments

First and foremost, we would like to thank the Government of the Philippines’ Department of Health and the Philippine Red Cross for supporting us with information on dengue and the country’s health system. We thank Steeve Ebener and Effie Espino for providing important inputs for the accessibility analyses. We are especially grateful to Kemal Arslantas from the Netherlands Red Cross, who initiated and led his organization’s efforts into understanding and quantifying epidemic risk in the past years; this work is also his legacy. Also from the Netherlands Red Cross, we would like to thank several researchers who contributed to the project: Annelot van Amerongen, Ayza Teng, Bart Veneman, Carla Meijerink, Chaima Abarkan, Dorike Jonker, Elena Stan, Elise Garton, Floor Lammers, Julia van den Berg, Lotte Schuitmaker, Merel van Cooten, Mruga Gurjar, and Tessa van Elsacker. Jacopo Margutti and Marc van den Homberg were financially supported by the Princess Margriet Fund. We thank the following students of the University of Geneva who have contributed to the development of this work and we would like to thank them: Alma Nurmuldina, Coralie Stavridis, Irène Daubard, Julie Seemann-Ricard, Maryam Cissé, and Quentin Pourrier.

